# Multiplexed effector screening for recognition by endogenous resistance genes using positive defense reporters in wheat protoplasts

**DOI:** 10.1101/2023.04.30.538885

**Authors:** Salome Wilson, Bayantes Dagvadorj, Rita Tam, Lydia Murphy, Sven Schulz-Kroenert, Nigel Heng, Emma Crean, Julian Greenwood, John P. Rathjen, Benjamin Schwessinger

## Abstract

- Plant resistance (*R*) and pathogen avirulence (*Avr*) gene interactions play a vital role in pathogen resistance. Efficient molecular screening tools for crops lack far behind their model organism counterparts, yet they are essential to rapidly identify agriculturally important molecular interactions that trigger host resistance.
- Here, we have developed a novel wheat protoplast assay that enables efficient screening of Avr/R interactions at scale. Our assay allows access to the extensive gene pool of phenotypically described *R* genes because it does not require the overexpression of cloned *R* genes. It is suitable for multiplexed *Avr* screening, with interactions tested in pools of up to fifty *Avr* candidates.
- We identified Avr/R-induced defense genes to create promoter-luciferase reporter. Then, we combined this with a dual-color ratiometric reporter system that normalizes read-outs accounting for experimental variability and Avr/R-induced cell-death. Moreover, we introduced a self-replicative plasmid reducing the amount of plasmid used in the assay.
- Our assay increases the throughput of *Avr* candidate screening, accelerating the study of cellular defense signaling and resistance gene identification in wheat. We anticipate that our assay will significantly accelerate *Avr* identification for many wheat pathogens, leading to improved genome-guided pathogen surveillance and breeding of disease-resistant crops.

## Introduction

Plant pathogenic fungi cause severe damage to cereal crops. These pathogens secrete hundreds of proteinaceous molecules called effectors during infection which manipulate plant processes to facilitate disease (Dodds & Rathjen, 2010). Some effectors can be recognized by resistance (R) proteins and are therefore also called avirulence (Avr) proteins. Avr recognition triggers a suite of defense-related host responses to fight off pathogen infection including the hypersensitive response (HR) that leads to cell-death at the site of infection and halts pathogen growth (Laflamme *et al*., 2020). The deployment of suitable *R* genes is a major breeding objective to reduce the negative impact of pathogens on crop production including in wheat. Recent years have made significant progress in identifying and cloning a wide variety of *R* genes in wheat (Wulff & Krattinger, 2022). Yet the identification of the corresponding *Avr* genes is lacking far behind. For example, there are only four confirmed *Blumeria graminis* s. str. (formerly known as *Blumeria graminis* f. sp*. tritici*) *Avr* genes (Praz *et al*., 2017; Bourras *et al*., 2019; Hewitt *et al*., 2021; Müller *et al*., 2022), three for *Zymoseptoria tritici* (Zhong *et al*., 2017; Meile *et al*., 2018; Amezrou *et al*., 2023), and five for *Puccinia graminis* f. sp. *tritici* (*Pgt*) (Chen *et al*., 2017; Salcedo *et al*., 2017; Upadhyaya *et al*., 2021; Arndell *et al*., 2023). There are no confirmed *Avr* genes for *Puccinia triticina* and *Puccunia striiformis* f. sp. *tritici* despite both belonging to the top five wheat pathogens globally (Savary *et al*., 2019). This lack of confirmed *Avr* genes in wheat pathogens and especially rust fungi is confounded by the absence of an efficient screening tool for *Avr* recognition in wheat. Most current methods require cloned *Avr* and *R* gene candidates that retain function when expressed heterologously and exhibit HR as general read-out for Avr/R interactions. For example, the model plant *Nicotiana benthamiana* is used to validate Avr/R interactions as it produces visible HR upon transient expression of *Avr/R* genes through agroinfiltration. Virus-mediated overexpression (VOX) can be used to screen *Avr* candidates in wheat cultivars carrying the corresponding *R* gene as the Avr recognition will lead to visible HR symptoms on the infected leaves (Lee *et al*., 2012; Bouton *et al*., 2018). However, the method is only applicable to small proteins (Lee *et al*., 2012) and used viruses are not accessible globally due to biosecurity regulations (Bouton *et al*., 2018). Recently, HR-based assays in wheat protoplasts have emerged as a potential alternative to test Avr/R interactions (Saur *et al*., 2019). In this transient assay, wheat protoplasts are co-transfected with *Avr/R* genes along with a *luciferase* reporter gene. Avr/R interaction triggers cell-death causing significant decreases in luciferase enzyme activity. This assay has been used to validate *Avr* candidates from wheat rust pathogens (Saur *et al*., 2019; Luo *et al*., 2021; Upadhyaya *et al*., 2021; Ortiz *et al*., 2022). However, the HR-based wheat protoplast assay in its current form has several limitations that make it impractical for high-throughput *Avr/R* screening. One such limitation is that it requires the transient overexpression of both the *Avr* and *R* genes from plasmids (Saur *et al*., 2019). This prevents screening against the large number of *R* genes which have been phenotypically characterized but not yet been cloned (McIntosh *et al*., 1995). Also, it is well known that luminescence is highly variable between biological replicates and experiments (Saur *et al*., 2019; González-Grandío *et al*., 2021) which can make the results difficult to interpret for high-throughput screening purposes. The assay in its current form relies on the reduction of luciferase activity as read out which is unspecific and cannot be differentiated from cell-death that is unrelated to HR. The assay in its current form also requires the preparation of large quantities of target gene plasmids for the protoplast transfection to achieve observable Avr/R-triggered cell-death (Saur *et al*., 2019; Upadhyaya *et al*., 2021; Ortiz *et al*., 2022). These limitations encouraged us to develop a novel assay in wheat protoplast that relies on a positive defense specific read-out and accounts for experimental variability via normalization, which is common practice for protoplast report assays (González-Grandío *et al*., 2021). Our new assay allows the research community to access the whole wheat gene pool to screen for Avr/R interactions and enables the investigation of downstream defense signaling directly in wheat.

## Results

### Profiling temporal HR onset induced by Avr/R interactions in wheat protoplasts

Our end goal was to identify defense signaling responsive promoter elements downstream of Avr/R recognition in wheat protoplast before or independent of the onset of HR. Hence, we first sought to characterize the time of HR onset in wheat protoplasts following Avr/R recognition to identify a suitable time point for RNAseq expression analyses. We used two published *Avr/R* gene pairs, *AvrSr35/Sr35* (Saintenac *et al*., 2013; Salcedo *et al*., 2017) and *AvrSr50/Sr50* (Mago *et al*., 2015; Chen *et al*., 2017), from wheat stem rust *P. graminis* f. sp. *tritici* for initial method development. As previously described (Saur *et al*., 2019), we used luminescence as a proxy for cell viability after overexpression of the *luciferase* reporter gene in conjunction with each *Avr/R* combination and corresponding control. When expressed from a plasmid, *Avr* and *R* genes were under control of maize ubiquitin promoter (*UBI*) (Christensen *et al*., 1992). We used the wheat cultivar (cv.) Fielder for the analysis of *AvrSr35/Sr35* and expressed both genes from a plasmid. For *AvrSr50/Sr50*, we made use of the introgression of *Sr50* into cv. Gabo (GaboSr50) and generated protoplast from cv. Gabo and cv. GaboSr50 (Jensen & Saunders, 2023). We only expressed *AvrSr50* from a plasmid. We also included the controls, the reporter alone and the reporter with *Avr* or *R* alone (Zenodo Supplemental Data S1). We measured luminescence at 3, 4, 5, 6, and 18 h post-transfection (hpt). We observed a gradual increase in luminescence measurements in protoplast expressing luciferase alone or when expressed together with an *Avr* or *R* gene only (Figure 1, A and B). The luciferase activity was strongest at 18 hpt in all treatment groups except for the ones expressing *Avr/R* pairs. In cv. Fielder expressing *AvrSr35*/*Sr35* and in cv. GaboSr50 expressing *AvrSr50*, luminescence measurements only increased slightly early on during the time course when compared to mock treatments of protoplasts transfected with no DNA (red line, Figure 1 C). At about 4 to 5 hpt we observed a divergence of luciferase activity from control treatment groups when compared with the Avr/R interaction treatment groups (Figure 1A and B). The lack of continuous increase of luciferase activity specifically in protoplast expressing *Avr*/*R* pairs suggests that their co-expression leads to cell-death in wheat protoplasts soon after transfection.

**Figure 1:**
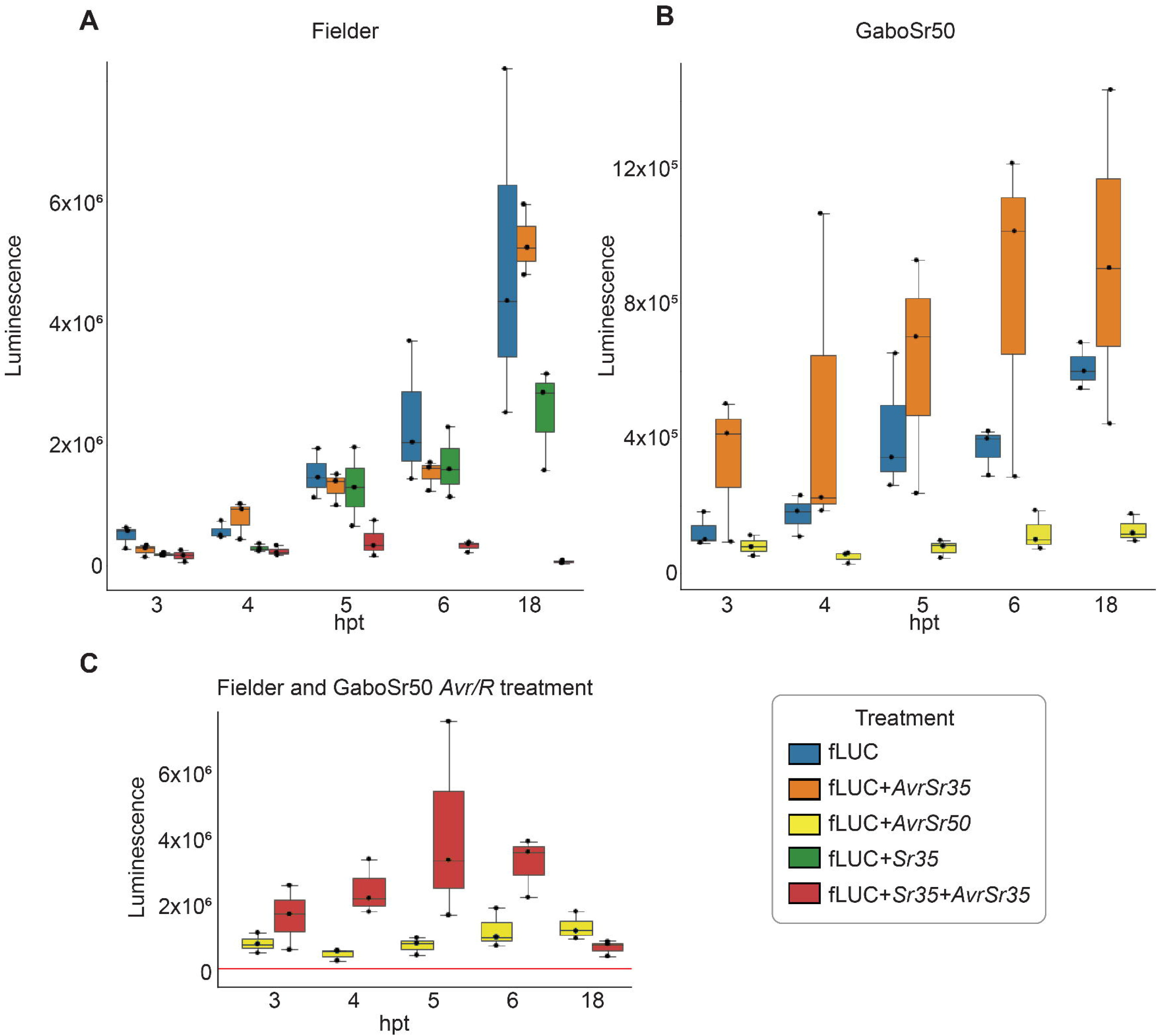
Rapid onset of cell-death downstream of *Avr/R* interactions in protoplasts. **A-C**, Luciferase activity was measured at defined timepoints in hours post-transfection (hpt) in wheat protoplasts. **(A)** Shows boxplots of cv. Fielder protoplasts expressing *luciferase* alone, and *luciferase* with either *Sr35, AvrSr35,* or *Sr35* plus *AvrSr35*. **(B)** shows boxplots of cv. GaboSr50 protoplasts expressing *luciferase* alone, and *luciferase* with either *AvrSr35* or *AvrSr50.* **(C)** shows the luminescence of the Avr/R interaction treatment groups from cv. Fielder and cv. GaboSr50 alone when compared to the no plasmid transfection control replicates (red line) which is the average of nine replicate across the six timepoints.

### Transfected protoplasts are suitable for replicable transcriptional analysis

We concluded that 4 hpt is a suitable time point to capture *Avr/R*-dependent transcriptional changes before strong HR induction based on our time course analysis (Figure 1). We repeated the experimental setup while omitting the reporter plasmid and collected protoplasts at 4 hpt for RNA sequencing (Figure 2 A, Zenodo Supplemental Data S2). We included four biological replicates per treatment group and sequenced each sample on the Illumina platform generating >20 million 150 bp paired-end reads per replicate. We used kallisto (Bray *et al*., 2016) for transcript quantification and DESeq2 (Love *et al*., 2014) for downstream differential expression analysis. As a first quality control step we quantified the expression of all *Avr* and *R* genes to investigate their specific expression pattern at 4 hpt. This analysis clearly showed that all genes are correctly expressed and at similar levels, including in treatment groups that co-express *Avr/R* gene pairs (Figure 2 A). In addition, it revealed that the endogenous expression of *Sr50* in cv. GaboSr50 is much lower level than the overexpressed of *Sr35* in cv. Fielder from a plasmid under the *UBI* promoter. Next, we performed principal component analysis (PCA) of all sample replicates combined. This revealed that the factor “wheat cultivar” had the strongest impact on the observed variation in transcript abundance and accounted for close to 90% of all variation on the first principal component (PC1), separating the two Gabo cultivars from cv. Fielder and further separating cv. Gabo from cv. GaboSr50 along principal component two (PC2) (Figure 2 B). Consequently, we analyzed each cultivar separately. The wheat cultivar specific PCA analysis showed a clear impact of Avr/R interaction on transcript abundance variation for both *AvrSr35/Sr35* and *AvrSr50/Sr50* (Figure 2 C and D). This impact was replicable because all four biological replicates clustered closely together and clearly separated treatment groups including Avr/R interactions sample types from their controls. This is consistent with the cell-death results seen in Figure 1 A and B. These results suggest that these data could be used to identify reporter genes specifically expressed downstream of Avr/R interactions in wheat protoplasts.

**Figure 2:**
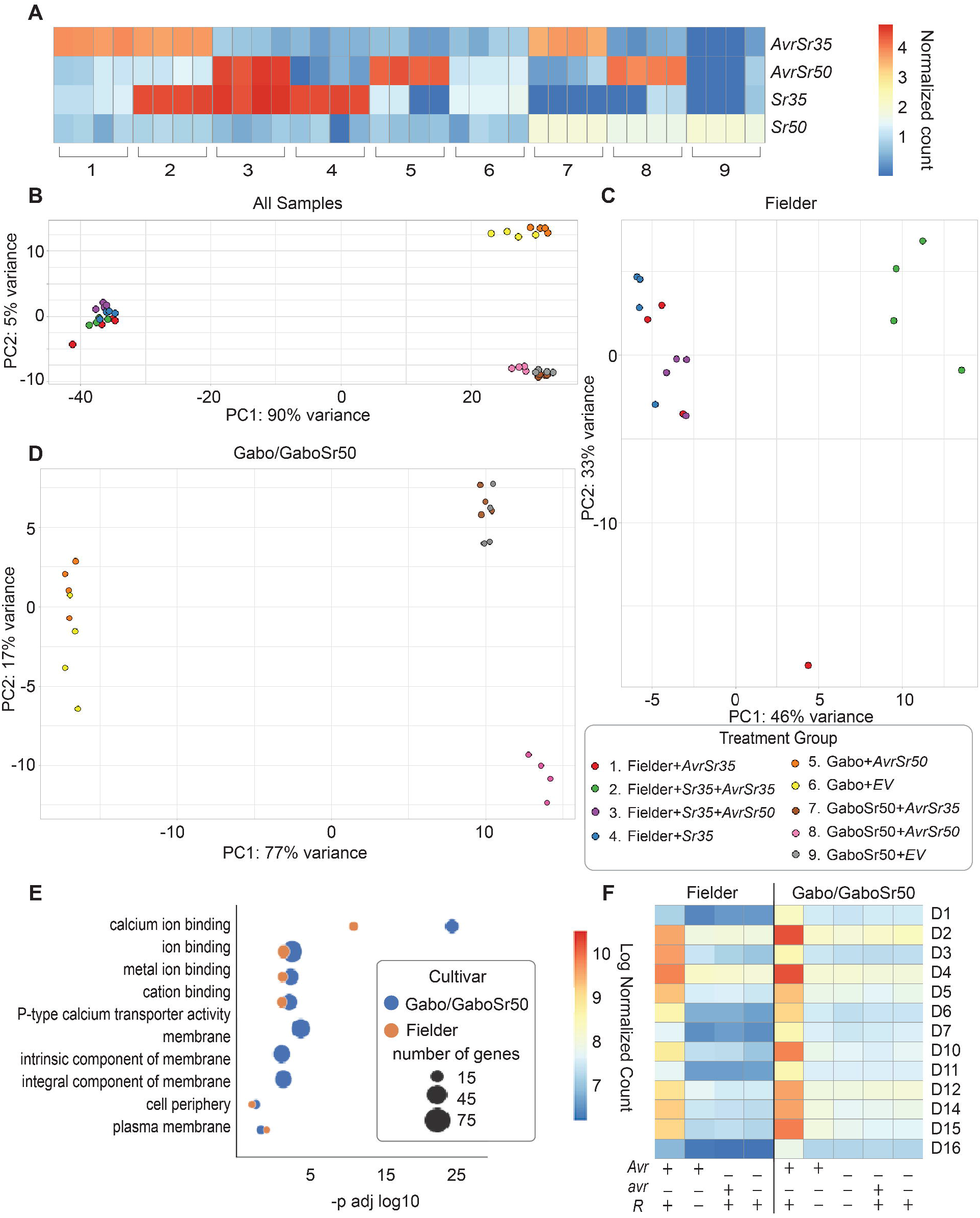
Co-expression of *Avr/R* gene pairs triggers defense signaling in wheat protoplasts. **(A)** Heat map shows expression of *Avr* and *R* genes by normalized gene count. Each column represents a replicate and the treatment group is indicated by a number as shown in the legend. *R* or *Avr* genes are expressed in their respective treatment groups only. This includes endogenously expressed *Sr50* in cv. GaboSr50. **(B-D)** show principal component analysis (PCA) of the RNAseq analysis; all 36 samples **(B)**, cv. Fielder **(C)** and cv. Gabo/GaboSr50 **(D)**. Figure legend numbers corresponds to **(A)**, while colors correspond to **(B), (C)** and **(D)**. (**E)** shows the GO terms of genes that are upregulated 2 fold in *Avr/R* treatment groups. A full list of upregulated genes with overrepresented GO terms is recorded in supplemental data (Zenodo Supplemental Data S4). **(F)** shows a heatmap of the expression of thirteen candidate reporter genes (D1-7, D10-12, D14-16) that are commonly upregulated by *Sr50/AvrSr50* and *Sr35/AvrSr35* in cv. GaboSr50 or cv. Fielder, respectively. Each square represents the mean of four replicates for treatment group.

### *Avr/R* recognition in wheat protoplasts leads to defense signaling

Next, we investigated if the observed gene expression changes in *Avr/R* expressing protoplasts overlapped with gene ontology (GO) terms commonly associated with defense signaling pathways in wheat and other plant species. We identified all genes that are specifically upregulated in response to Avr/R interactions in cv. Gabo/Gabo50 and cv. Fielder above a 2-fold change threshold. We identified 309 and 922 genes in *AvrSr35/Sr35* and *AvrSr50/Sr50* expressing protoplasts, respectively, when compared to all their control treatment groups. Complete gene lists are available on Zenodo as Supplemental Data S3. We performed a statistical enrichment analysis using these gene lists to test for overrepresentation of defense signaling associated GO terms. Our results revealed significant enrichment for GO terms related to cellular components of cell periphery (GO:0071944) and plasma membrane (GO:0005886) in both cultivars (Figure 2 E). We identified defense response to other organisms (GO:0098542), response to other organisms (GO:0051707), and biological process involved in interspecies interaction between organisms (GO:0044419) when looking at cv. GaboSr50 alone (Zenodo Supplemental Data S4). We found molecular functions GO terms being enriched for calcium ion binding (GO:0005509), cation binding (GO:0043169), and calcium ion transmembrane transporter activity (GO:0015085) in both cultivars (Figure 2 E). Our GO terms analysis suggests that *Avr/R* expressing protoplasts undergo significant transcriptional reprogramming consistent with calcium-mediated defense signaling before the onset of strong cell-death.

We were then interested in genes that are commonly upregulated by the Avr/R interaction in both cv. GaboSr50 and cv. Fielder. We identified 66 genes commonly upregulated in both cultivars, (Zenodo Supplemental Data S5) of which only 16 genes were associated with three different GO terms, all related to the molecular function of ion binding (Zenodo Supplemental Data S4). Our results showed that several of the identified genes have been previously implicated in plant defense signaling including *TaNHL10* (*TRAESCS3D02G368800*) (Century *et al*., 1995; Aarts *et al*., 1998; Dagvadorj *et al*., 2022).

In summary, we observed the upregulation of genes involved in defense signaling in wheat protoplasts expressing *Avr/R* pairs. The promoters of commonly upregulated genes, by two *Avr/R* pairs and in two wheat cultivars (Figure 2F), are promising candidates to generate Avr/R-inducible reporters.

### Identification of positive defense reporters in wheat protoplasts

We further investigated the expression patterns of the 66 commonly upregulated genes to select a subset whose promoters we could test as positive defense signaling reporters. We selected 13 promoters (D1-D7, D10-D12, and D14-D16) based on their genes’ expression patterns in both cultivars (Figure 2 F). To further evaluate these candidates, we cloned or synthesized their promoters, consisting of 600-800 bp upstream of their respective genes’ start codons. We accounted for the impact of Avr/R-induced HR upon luciferase activity (Figure 1) by adapting a dual-color green/red luciferase reporter system (González-Grandío *et al*., 2021). This ratiometric system uses green- and red-shifted luciferases (Sarrion-Perdigones *et al*., 2019), where green-shifted luciferase is fused to a conditionally responsive promoter and red-shifted luciferase serves as a constitutively expressed reference for internal normalization, thus greatly reducing intra experimental variation.

To test the activity of our thirteen candidate defense promoters, we generated promoter fusion constructs driving the expression of green-emitting luciferase (E-Luc). These plasmids are named *pDefense* (*pD*[*1-7, 10-12, 14-16*]), respectively. We co-transfected these plasmids along with internal normalization plasmid *UBI-pRedf* and *UBI-AvrSr50* (Supplemental Figure 1) into protoplasts isolated from cv. GaboSr50 and Gabo. For the initial activity screening, we performed the test with two technical replicates. Ten of these promoters did not result in specific and reproducible promoter activation downstream of AvrSr50/Sr50 interactions when compared to their respective controls. In contrast, the three promoters D2, D14, and D15 showed a strong increase in normalized ratiometric luminescence specifically downstream of AvrSr50/Sr50 interactions (Supplemental Figure 2). Figure 3 shows an example of the specific activation of D14 downstream of AvrSr50/Sr50 in cv. GaboSr50 when compared to controls not expressing the *R* gene or the matching *Avr*. These results indicate that our new ratiometric reporter system is able to detect specific defense activation downstream the Avr/R interactions when the *R* gene is expressed endogenously.

**Figure 3:**
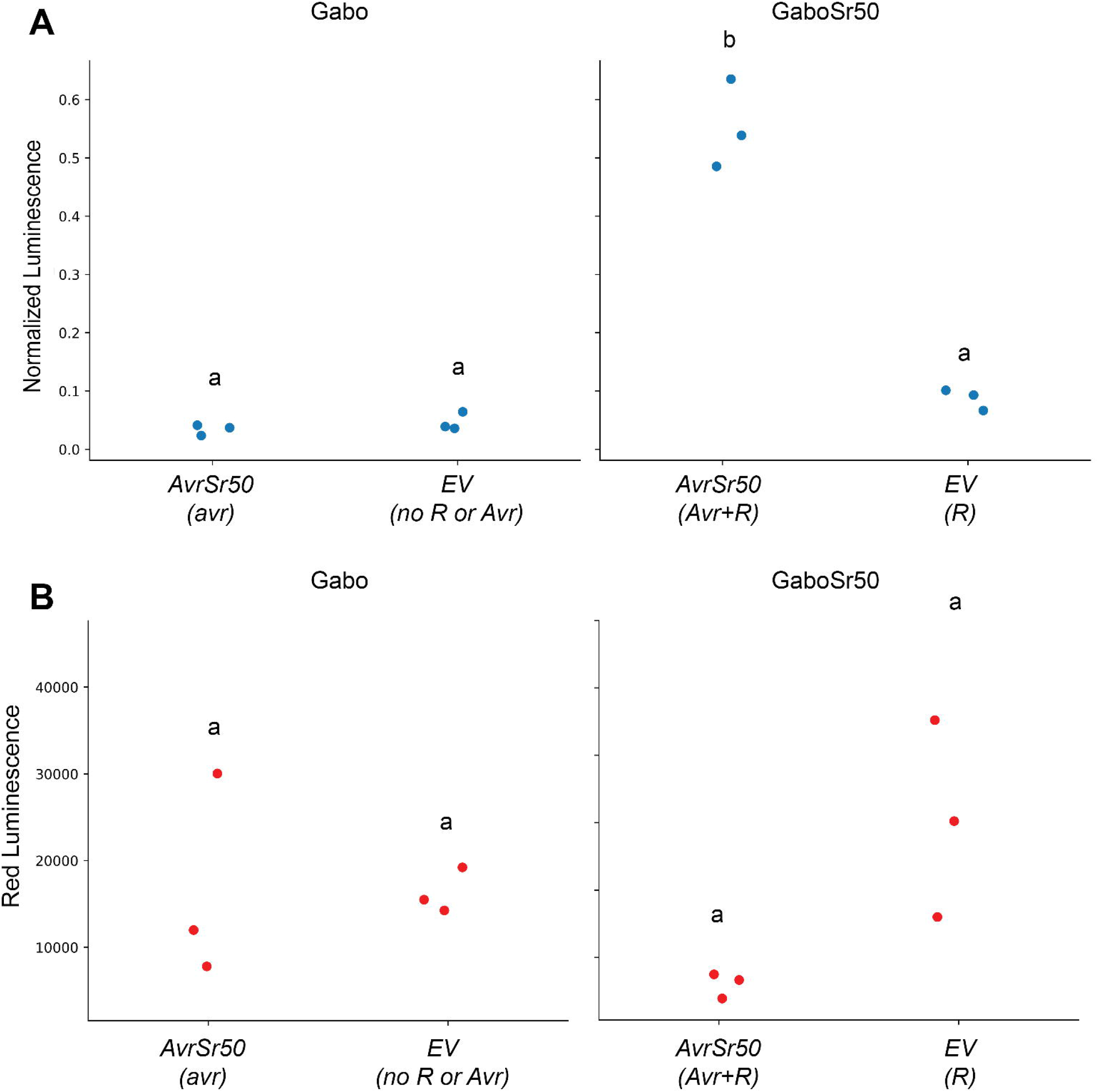
*pDefense14* is a positive defense reporter in wheat protoplasts. (**A**) Wheat protoplasts transfected with *pDefense14* and *UBI-pRedF* constructs show an increase in normalized luminescence in the *Avr/R* treatment, compared to treatments with *R* or *avr* only and without *R* or *Avr*. Panel **(B)** shows corresponding non-normalized red-filtered luminescence for the treatments in panel A. Treatments marked with common letters were not significantly different (P >0.05) assessed with one-way analysis of variance (ANOVA) and post hoc Tukey’s HSD test.

### The wheat dwarf virus-derived replicons-based system further increases the sensitivity of the positive defense reporter assay

The current wheat protoplast assay requires a large amount of DNA material for transfection to obtain observable cell-death. This makes its application prohibitive for large scale screening of many candidates or variants. We tested a replicon-based system using a disarmed version of the wheat dwarf virus (WDV) in our expression vector backbone (Gil-Humanes *et al*., 2017). We reasoned that this would generate self-replicating plasmids in wheat protoplasts and thereby reduce the amount of plasmids required to detect defense activation in our new ratiometric wheat protoplast assay. We assembled Golden Gate compatible WDV vector from Gil-Humanes et al. (2017) and denoted it as *pWDV1* (see Material and Methods). We expressed *AvrSr50* under ubiquitin promoter using *pWDV1* (*pWDV1-AvrSr50*) and used it to generate dose response curves using different amounts (0.1, 0.5, 1, 5 and 15 µg) of plasmid DNA for transfection. We measured induction of D14 induction (normalized luminescence) and cell viability (red luminescence) and compared it to the dose response of the standard non-viral plasmid *pICH4772*-*AvrSr50* (Figure 4). We observed specific D14 promoter induction in all samples of cv. GaboSr50 protoplasts transfected with *pICH4772*-*AvrSr50* and *pWDV1-AvrSr50* when compared to controls in cv. GaboSr50 transfected with *pWDV1-AvrSr35* (1 and 15 µg) and cv. Gabo transfected with *pWDV1-AvrSr50* (1 µg) or *pWDV1-AvrSr35* (1 µg). We also observed that the promoter induction increases relative to the amount of *pICH4772*-*AvrSr50* used. At the same time, we were still able to observe induction of the defense reporter D14 even in the absence of cell-death at 0.1, 0.5 and 1 µg plasmid used. Using *pWDV1-AvrSr50* further improved the sensitivity of the assay as it increased defense reporter D14 activation at 0.1 and 0.5 µg plasmid and decreased cell viability when compared to the same amounts of plasmid for *pICH4772*-*AvrSr50*. Altogether, this result indicates that the ratiometric dual-color luciferase system combined with *pDefense* reporter and *pWDV1* is highly sensitive and specific requiring much less plasmid to detect Avr/R interactions than a single luciferase-based wheat protoplast assays.

**Figure 4:**
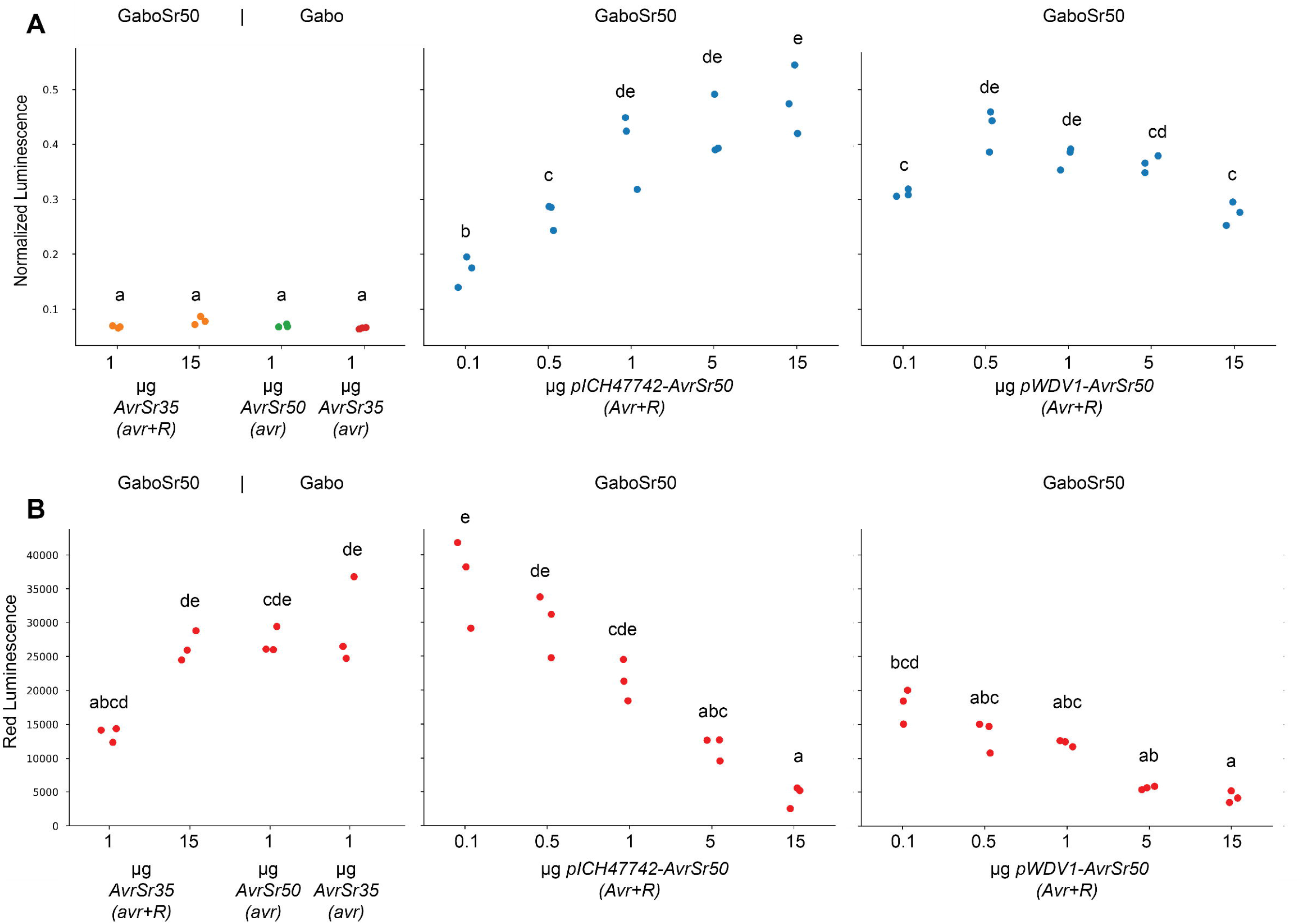
*pWDV1* plasmid reduces amount of *Avr* plasmid required to detect defense signaling using *pDefense* reporter. *AvrSr50* was cloned into *pWDV1* (*pWDV1-AvrSr50*) and non-viral plasmid backbone (*pICH4772*-*UBI-AvrSr50)* and transfected in increasing amounts (0.1, 0.5, 1, 5 and 15 µg of plasmid DNA) alongside *pDefense14* and *UBI-pRedf.* Reporters remained in the non-replicative plasmid. Controls in cv. GaboSr50 were transfected with *pWDV1-AvrSr35* (1 and 15 µg) and cv. Gabo transfected with *pWDV1-AvrSr50* (1 µg) or *pWDV1-AvrSr35* (1 µg). One-way analysis of variance (ANOVA) and post hoc Tukey’s HSD test was used to assess differences within normalized luminescence **(A)** and red luminescence **(B)** groups. Treatments marked with a common letter were not significantly different (P >0.05).

### Positive defense reporters are highly sensitive to detect intracellular Avr/R interactions in wheat protoplast expressing *R* genes endogenously

Next, we explored if the defense reporters require overexpression of the *R* gene and if they are applicable to additional Avr/R interactions. We tested the induction of D14 expressing *AvrSr27, AvrSr35,* and *AvrSr50* in *pWDV1* using 1 µg plasmid, instead of the previously used 15 µg. We transfected cv. Coorong, C6969, and GaboSr50, with *pWDV1* encoding the recognized *Avr*, an unrecognized or non-matching *avr*, or the empty vector control (Figure 5). Wheat cv. Coorong carries *Sr27* (Upadhyaya *et al*., 2021), and cv. C6969 (*Triticum aestivum*) carries *Sr35* which was introgressed from *Triticum monococcum* (McIntosh *et al*., 1984). We observed a strong induction of D14 specifically downstream of AvrSr27/Sr27, AvrSr35/Sr35, and AvrSr50/Sr50 interactions but not in any of the controls (Figure 5 A-C), validating the D14 defense promoter as well as *pWDV1* applicability in wheat protoplast assay. We decided to focus on D14 and switch all *Avr* expressions in *pWDV1* for the remainder of the study but to keep the reporter genes in the non-viral vector. We further tested the D14 promoter with two newly reported stem rust Avr proteins AvrSr13 and AvrSr22 (Arndell *et al*., 2023) by transfecting them in wheat cv. Line S and W3534 (Park, 2016), respectively (Figure 5 D and E). Similar to the other *Pgt* AvrSr/Sr interactions, we observed strong specific induction of D14 from AvrSr13/Sr13 and AvrSr22/Sr22 combinations.

**Figure 5:**
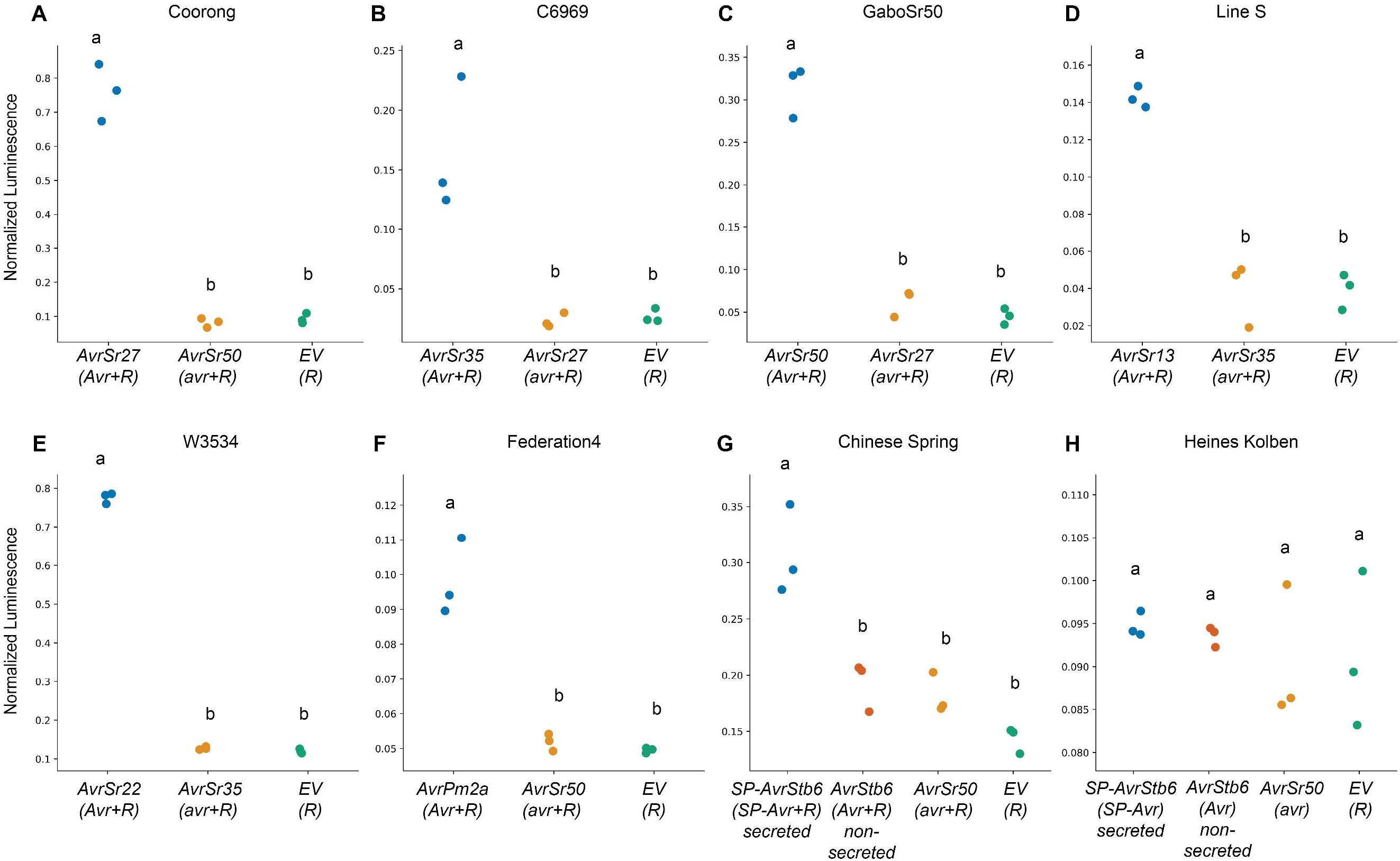
Positive *pDefense* reporter detects defense signaling triggered by seven distinct *Avr/R* pairs in wheat protoplasts expressing the *R* gene endogenously. Protoplasts from wheat cultivars **(A)** cv. Coorong carrying *Sr27*, **(B)** cv. C6969 carrying *Sr35,* (C) cv. GaboSr50 carrying *Sr50,* (D) cv. Line S carrying *Sr13*, **(E)** cv. W3534 carrying *Sr22*, **(F)** cv. Federation4/Ulka carrying *Pm2* and **(G)** cv. Chinese Spring carrying *Stb6* were transfected with *UBI-pDefense14* and *Avr* constructs, as well as an *avr* (unrecognized *Avr*) construct and empty vector **(EV)** construct as controls. **(H)** had identical treatments to **(G)**, however cv. Heines Kolben lacks *Stb6. UBI-pRedf* was used for normalization of all experiments. For **(H)** and (G) *AvrStb6* was expressed with (SP*-AvrStb6)* and without (*AvrStb6*) secretion peptide to allow for extracellular recognition including the appropriate control. One-way analysis of variance (ANOVA) and post hoc Tukey’s HSD test was used to assess differences among normalized luminescence for each cultivar. Treatments marked with a common letter were not significantly different (P >0.05).

To assess the D14 promoter specificity to Avr/R interaction, we exploited the natural and mutational variation of *AvrSr50* alleles that mediate the recognition by *Sr50* (Ortiz *et al*., 2022). We tested three AvrSr50 variants displaying varying interaction strength with Sr50; avrsr50-A1 which carries a single amino acid substitution at Q121K and is not recognized by Sr50, avrsr50-B6 which is found in *Pgt* race QCMJC and is not recognized by Sr50 (Chen *et al*., 2017) and AvrSr50-C, which is found in *Pgt* race RKQQCs and is recognized by Sr50 (Chen *et al*., 2017) in cv. GaboSr50 (Supplemental Figure 3). The previously used *AvrSr50*, *AvrSr35* and empty vector served as controls. Using our assay, we confirmed previous results that the specific single amino acid substitutions in avrsr50-A1 and avrsr50-B6 were sufficient to abolish recognition by Sr50, while changes to six different amino acids in AvrSr50-C only partially abrogated recognition by Sr50 (Supplemental Figure 3). This demonstrates that the D14 promoter successfully detects specific Avr/R interaction in allelic variants of Avr proteins such as AvrSr50.

Next, to show the versatility of D14 promoter, we tested AvrPm2 from *Blumeria graminis* s. str. (Praz *et al*., 2017) (Figure 5F). We observed specific D14 induction compared to the controls when we transfected *AvrPm2* into wheat cv. Federation4/Ulka, which carries *Pm2* (Sánchez-Martín *et al*., 2021).

These results support that the D14 promoter in combination with *pWDV1* is broadly applicable as a defense reporter in wheat independent of the wheat cultivar and the specific *Avr/R* combination used. This suggests that our protoplast assay is applicable to testing wheat cultivars that carry endogenous *R* genes and does not require a cloned *R* gene.

### Positive defense reporter D14 is able to detect extracellular AvrStb6/Stb6 interaction in wheat protoplast

We were interested in the induction of D14 reporter by extracellular Avr/R interactions. For this purpose, we tested AvrStb6 from *Zymoseptoria tritici* (Zhong *et al*., 2017), which is recognized by *Stb6* encoding a wall-associated receptor kinase-like protein (Saintenac *et al*., 2018). We decided to express AvrStb6 including its signal peptide (SP) allowing for AvrStb6 accumulation outside the cells to allow for extracellular recognition of AvrStb6 by Stb6. We included the non-secreted form of AvrStb6 without SP as negative control. We observed significant D14 induction compared to the controls when we transfected secreted-AvrStb6 in protoplasts derived from wheat cv. Chinese Spring which carries *Stb6* (Supplemental Figure 4) (Saintenac *et al*., 2018). The induction of D14 was specific to wheat lines carrying *Stb6* because we did not observe it in wheat cv. Heines Kolben which lacks *Stb6* based on our PCR screening (Supplemental Figure 4). This result indicates that our defense reporter assay is able to detect extracellular Avr/R interactions and validates versatility of the assay among Avr proteins from different wheat pathogens.

### Positive defense reporters are broadly applicable for Avr/R interaction screening in many wheat cultivars

We aimed to illustrate the broad applicability of our new assay in a wide range of wheat cultivars as current cell-death based assays were mostly conducted in cv. Fielder (Saur *et al*., 2019, p. 5; Luo *et al*., 2021; Upadhyaya *et al*., 2021; Ortiz *et al*., 2022; Arndell *et al*., 2023). We tested the defense signaling reporter D14 for its inducibility by three *Avr/R* combinations in protoplasts of nine different wheat cultivars. We tested cultivars that carry *R* genes (cv. GaboSr50, cv. Coorong and cv. C6969) with their corresponding *Avr* construct *(AvrSr50, AvrSr27* and *AvrSr35,* respectively). The cultivars Clement, Compair, Avocet Yr8, (Dracatos *et al*., 2016) and Kenya W1483 (McIntosh *et al*., 1995) were tested with overexpression of *AvrSr50/Sr50* combination. The cultivars HeinesPeko (Dracatos *et al*., 2016) and Fielder were tested with *AvrSr27/Sr27* and *AvrSr35/Sr35* combinations, respectively. We included a negative control in all cultivars by expressing an unrecognized (non-matching to R protein) *avr, Avr* or *R* gene alone (Figure 6). In all cultivars we tested, a consistent robust induction of normalized luminescence of *pD14* was observed in protoplasts expressing cognate *Avr/R*-pair compared to the negative controls. The relative normalized luminescence and its induction in the *Avr/R* treatment group varied between the different cultivars. These results indicate that the positive defense reporter assay enables the detection of Avr/R interactions in many wheat cultivars.

**Figure 6:**
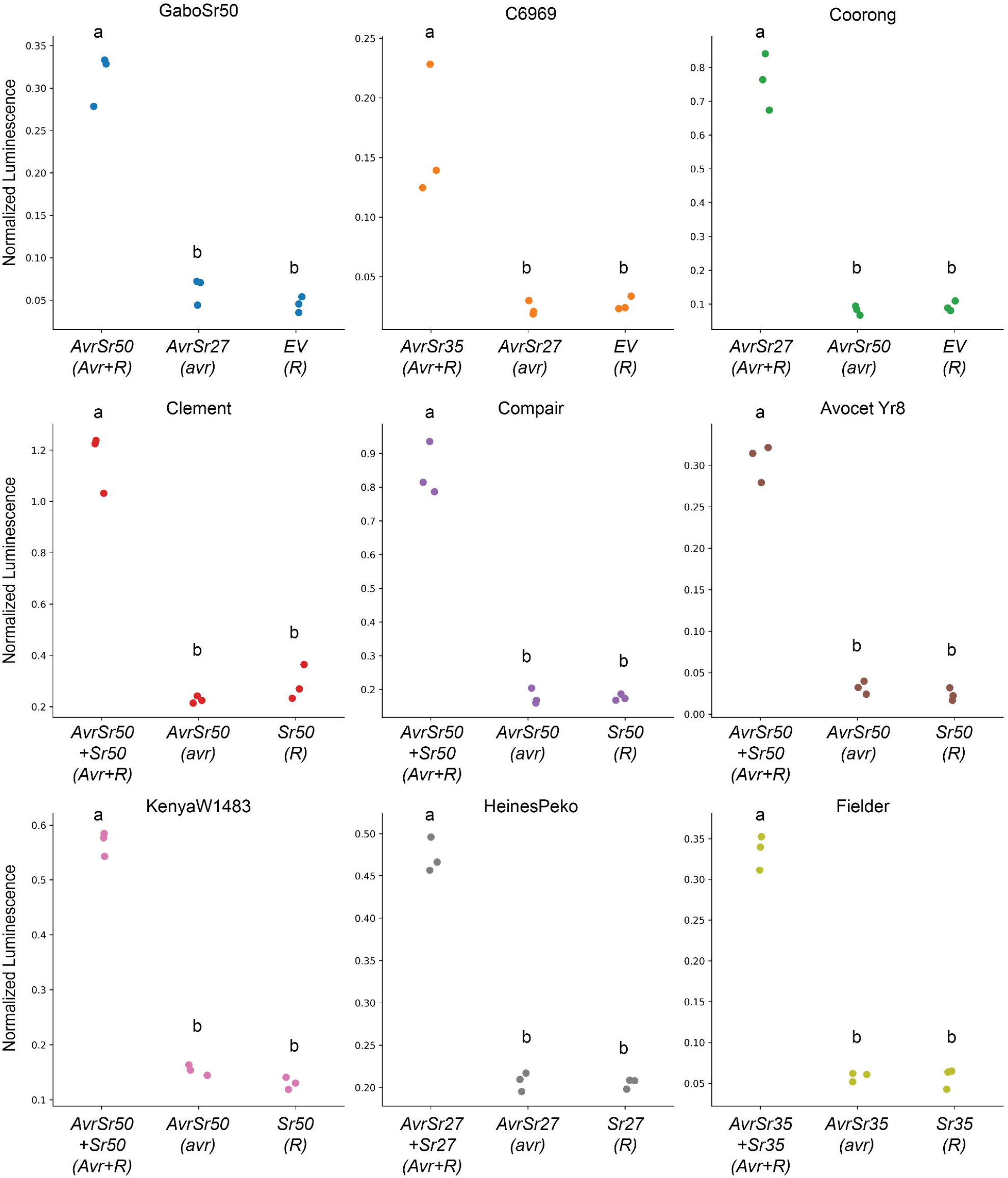
The positive defense reporter *pDefence14* is broadly applicable to all tested wheat cultivars. Each panel shows the normalized luminescence of the positive defense reporter *pDefense14*. Protoplasts isolated from various wheat cultivars were transfected with both *R* and *Avr* constructs, aside from cultivars GaboSr50, C6969 and Coorong which carry *Sr50, Sr35* and *Sr27* respectively. The cultivars Corack, Clement, Kenya W1483, Avocet Yr8 and Compair, tested with *AvrSr50/Sr50* and Heines Peko was tested with *AvrSr27/Sr27.* An *avr* (unrecognised by R protein) was used as a control and *UBI-pRedf* was used for normalization of all experiments. One-way analysis of variance (ANOVA) and post hoc Tukey’s HSD test was used to assess differences among means of normalized luminescence measurements for each cultivar. Treatments marked with common letters were not significantly different (P >0.05).

### The positive defense reporter assay enables multiplexed screening of *Avr* candidates in bulk

We aimed to test if our assay could be used for multiplexed screening of Avr/R interactions using AvrSr50/Sr50 as a proof of concept. We co-transfected a pool of twenty-four *Avr* candidates (Methods for details) into protoplasts isolated from cv. GaboSr50 and cv. Gabo in the presence or absence of *AvrsSr50*. The pools include 1 µg of each *pWDV1-Avr*. Each replicate included *pD14* reporter and *UBI-pRedf* co-transfection to detect *Avr/R*-triggered defense promoter induction and normalized luminescence measurements. In cv. GaboSr50 protoplasts, *AvrSr50* alone and the pooled mixes containing *AvrSr50* induced a high level of normalized luminescence of *pD14* when compared to the pooled mix without *AvrSr50* and all other treatment groups including controls in cv. Gabo protoplasts (Figure 7). We observed a clear specific induction of the D14 promoter only in the presence of *AvrSr50* in cv. GaboSr50 when provided as pairs or as multiplexed mixtures with no significant statistical difference between the single or multiplexed treatment groups (Figure 7). Overall, these results show that our positive defense reporter protoplast assay can be used to screen a pool of *Avr* candidates against a wheat cultivar endogenously expressing the recognizing *R* gene, which then can be followed up to identify the specific positive *Avr/R* interaction.

**Figure 7:**
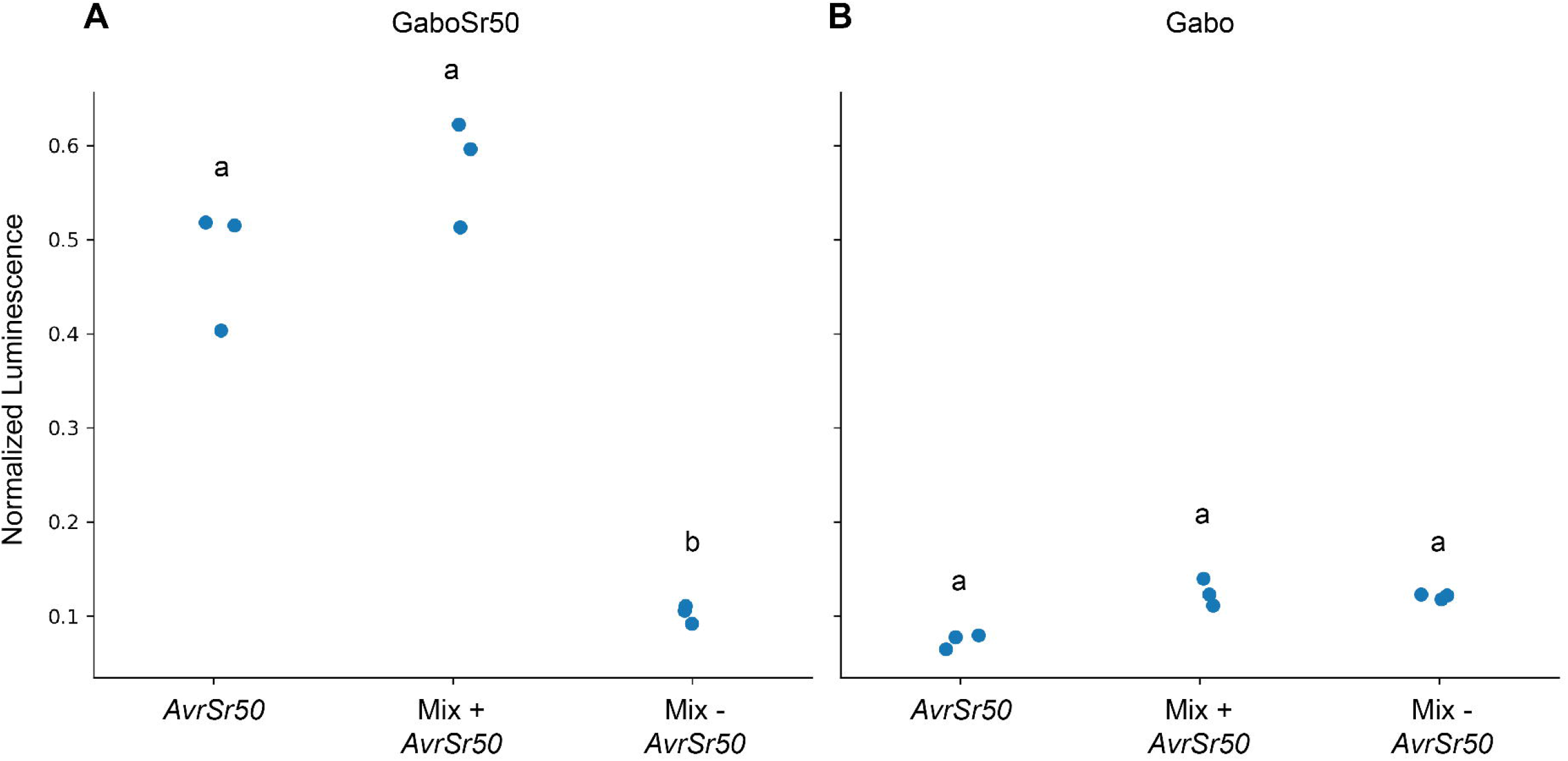
The positive defense reporter assay enables multiplexed *Avr* screening. *Avr* pools of twenty-five candidates were transfected into wheat protoplasts isolated from **(A)** cv. GaboSr50 and **(B)** cv. Gabo, along with *pD14* reporter and *UBI-pRedf*. *Avr* candidates included were; *AvrSr13, AvrSr22, AvrSr27, AvrSr35, AvrSr50, AvrStb6, Avr3D1, AvrStb9, AvrPm1, AvrPm2, AvrPm3 AvrPm17* and thirteen other wheat rust effector candidates. One µg of each *pWDV1-Avr* was used, with the negative control excluding *AvrSr50* and positive control of 1 µg of *AvrSr50* alone. For each cultivar, one-way analysis of variance (ANOVA) and post hoc Tukey’s HSD test was used to assess differences among means of normalized luminescence and red luminescence measurements for each treatment group. Treatments marked with common letters were not significantly different (P >0.05).

## Discussion

Here, we describe an improved wheat protoplast assay to screen avirulence effector candidates and study cellular defense signaling in wheat. We identify Avr/R-induced promoters and use them to generate reporter constructs that produce a positive readout during Avr/R interaction. We also reduce variability of luminescence measurement by adapting a ratiometric dual-color luciferase reporter system (Sarrion-Perdigones *et al*., 2019) and enable assays with candidate *Avr* plasmids prepared from simple minipreps. We show the broad applicability of the positive defense reporters by testing seven *Avr/R* pairs from three different wheat pathogens on thirteen different wheat cultivars. The improved wheat protoplast assay is highly sensitive because it allows for multiplexed screening of *Avr* candidates in wheat cultivars expressing *R* genes endogenously.

This approach has several advantages over previous methods applied to identify Avr/R interactions in wheat. First, it does not require a cloned and overexpressed *R* gene to test the interaction. We demonstrate that the positive defense reporter assay is suitable to test interactions directly in a broad range of wheat cultivars encoding specific *R* genes in their genome. This creates opportunities to screen *Avr* candidates directly on resistant wheat cultivars, which has been not easily possible with the cell-death based protoplast assays due to the inherent assay variability without normalization (Saur *et al*., 2019). Second, combining generic Avr/R-inducible promoters with the dual-reporter system generates normalized positive signals, which enables a much more sensitive screening approach by greatly reducing variability between replicates. Third, the assay uses 10-20 times less plasmid per *Avr* and screens multiple candidates in a single transfection, simplifying plasmid preparation and saving time and resources. Lastly, our new wheat protoplast reporter assay enables us to study a wide range of plant defense signaling directly in wheat. This is highlighted by our RNAseq analysis of defense signaling in wheat protoplasts. Our results show that genes responsible for calcium ion binding, cation binding and calcium ion transmembrane transporter activity along with other defense-related genes are specifically upregulated in response to both AvrSr35/Sr35 and AvrSr50/Sr50 interactions in two wheat cultivars. In line with our findings, there have been strong indications that plant defense signaling triggered by Avr/R interactions involves plasma membrane reorganization including formation of ion channels that regulate plant cell-death and disease resistance. Recent studies of *Arabidopsis* ZAR1 (Bi *et al*., 2021) and wheat Sr35 (Förderer *et al*., 2022) showed the activated ZAR1 and Sr35 proteins forming pentameric structures, so-called resistosomes, that appear to function as non-selective calcium channels responsible for Ca^2+^ uptake, leading to cell-death. Similarly, in *Arabidopsis*, an activated NRG1.1 protein, known as a downstream signaling component of plant *R* proteins, composes a Ca^2+^ permeable non-selective cation channel enabled by oligomerization, which controls cell Ca^2+^ influx and induces plant cell-death (Jacob *et al*., 2021). Therefore, our findings support that the channel activity, especially Ca^2+^ regulating channels, are conserved in broader Avr/R*-*induced downstream signaling.

Interestingly, one of our marker genes (the gene of promoter D14) encodes a predicted wheat NDR1/HIN1-like 10 (NHL10) protein. The NHL proteins that are grouped in a large gene family in *Arabidopsis* are involved in plant defense, such as disease resistance (Century *et al*., 1995; Aarts *et al*., 1998) and pathogen sensing (Zheng *et al*., 2004; Zipfel *et al*., 2004; Boudsocq *et al*., 2010). The NHL proteins have one or two transmembrane domain(s) (Zheng *et al*., 2004) and several of them were shown to localize to the plasma membrane (Coppinger *et al*., 2004; Selote *et al*., 2014; Dagvadorj *et al*., 2022), which is thought to be crucial for their functioning in disease resistance pathways. Moreover, *Arabidopsis NHL10* expression, which is highly induced by flg22, is reported to be involved in Ca^2+^ signaling through the activation of calcium-dependent protein kinases (Boudsocq *et al*., 2010). In a recent study, wheat blotch disease effector ToxA was shown to interact with a wheat NHL protein, TaNHL10, to promote cell-death mediated by intracellular resistance like protein Tsn1 (Dagvadorj *et al*., 2022). Future studies will benefit from our defense specific promoters, in particular D14, to investigate downstream defense signaling components in wheat.

Effectors are known to manipulate plant processes or suppress plant defense (Win *et al*., 2012) which could present challenges for screening large effector pools of unknown function. Some effectors may interfere with the defense signaling process resulting in inhibition of the defense promoter induction, which could prevent identifying a potential *Avr* candidate in the pool. This problem can be solved by including a known *Avr* as positive control and transfecting the pool into wheat cv. expressing cognate *R* gene of the known *Avr*. This positive control would reveal whether the pool having *Avr* candidate(s) interferes with the defense promoter induction. In addition, a split-pool approach that screens a limited number of subpools of *Avr* candidates with partial overlaps can further reduce the risk of specific suppression of a recognized *Avr* by another effector.

Future studies will further investigate the capabilities of our positive defense reporter based Avr screening assay. We have performed screening of twenty-five different *Avr* candidates in this study and we anticipate that the high sensitivity of the assay combined with self-replicative *pWDV1* plasmid can successfully screen far greater numbers of *Avr* candidates in a single wheat protoplast transfection. Moreover, we anticipate that the *pWDV1* plasmid will simplify transient gene overexpression in wheat protoplasts because it only requires miniprep plasmid samples (Supplemental Figure 5), avoiding labor intensive and expensive maxiprep plasmid purification. This will facilitate rapid functional gene and variant analysis directly in wheat as demonstrated for *AvrSr50* variants in our study.

Our assay can be combined with a recent report that describes highly multiplexed avirulence effector recognition screening using cell-death as a negative enrichment method (Arndell *et al*., 2023). Multiplexed effector libraries (n>600) of *Pgt* were co-transfected into cv. Fielder protoplasts overexpressing specific *Sr* genes from plasmids. The author used the overexpression of YFP (Yellow Fluorescent Protein) as a cell viability marker to separate alive from dead protoplasts via flow cytometry. This allowed for negative enrichment analysis of expressed effectors when comparing live protoplasts before and after flow cytometry via RNAseq analysis. The author could identify new matching *Avr* genes for two out of five *Sr* genes including *AvrSr13/Sr13* and *AvrSr22/Sr22* but not for *Sr21*, *Sr26*, and *Sr61*. Such large scale avirulence effector screening approaches will be much more efficient when combined with our positive reporter assays that will allow researchers to first identify avirulence effector pools that are recognized by a specific wheat cultivar. Further refinement of the reported negative enrichment screen will likely enable screening in wheat cultivars expressing *R* genes endogenously, similar to our assay. Application of both multiplexed avirulence effector screening approaches promises to usher in a transformative era of avirulence gene identification in economically important pathogens of wheat and other crop species.

## Methods

### Protoplast Isolation

Protoplast isolation and plant growth protocol are described here: https://dx.doi.org/10.17504/protocols.io.q26g7r3zkvwz/v1. Wheat plants of cv. Fielder, Gabo and GaboSr50 were grown for 7-9 days in a growth cabinet with 150 µmol/m^2^/s light intensity and at 21 °C with 16 hr day length. Protoplasts were isolated by making a shallow cut across the leaf epidermis and peeling back the epidermal layer to expose the mesophyll layer beneath. The cut segments were placed, peeled side down in 0.6 M Mannitol for 10 mins, then placed into a 7 cm diameter petri dish containing filtered enzyme solution (MES-pH5.7 (20 mM), Mannitol (0.6 M), KCI (10 mM), Cellulase (Yakult R-10) 1.5% w/v, Macerozyme (Yakult R-10) 0.75% w/v, CaCl_2_(10 mM). BSA 0.1% w/v). The dish was wrapped in aluminum foil to shield it from light and placed on an orbital shaker, horizontally rotating at 60 RPM, for three hours. Protoplasts were collected by gentle centrifugation at 100 x g for 3 mins in a 30 mL round-bottomed tube, then the enzyme solution was removed and the pellet was washed in W5 solution (MES-pH5.7 (2 mM), KCI (5 mM), CaCl_2_ (125 mM), NaCl (154 mM). Five µL of concentrated protoplasts were removed and used to quantify cells and check quality using a hemocytometer and light microscope. Protoplasts were washed again with W5, this time left on ice for 45 minutes to settle. W5 solution was removed, and protoplasts were resuspended to 300,000-350,000 protoplasts/mL using MMG (MES-pH5.7 (4 mM), Mannitol (0.4 mM), MgCl_2_ (15 mM)). Protoplasts were used immediately for transfection.

### Protoplast transfection

High concentration (1 µg/µL) plasmid DNA was extracted via maxi-prep (Promega SV Wizard). Lower quality plasmid DNA (e.g. from miniprep kits) typically resulted in low transfection rates. The *Avr*, *R* and *luciferase* genes were expressed under control of maize ubiquitin promoter (*UBI*), and 10 µg of each *Avr* and *R* plasmid and 20 µg of *luciferase* reporter plasmid were used for each replicate. Due to high concentration and purity, plasmids were heated at 65°C for 10 min before use to reduce viscosity and ensure consistent measurement and delivery into cells. The required plasmid DNA was added to 14 mL round-bottom culture tubes (Thermofisher, #150268). 300 uL of protoplasts and 300 µL of PEG solution (PEG 4000 (40% w/v), Mannitol (0.2 M), CaCl_2_ (100 mM)) was then added and mixed immediately by gently flicking/rotating the tube three times. Samples were incubated for 15 minutes, and the reaction halted with 900 µL W5. Protoplasts were then collected by 100 x g centrifugation for 2 min, resuspended in 500 µL modified W5 (W5 + 1 mM glucose) and transferred to a transparent, sterile 12 well culture plate (Costar, #3513) whose wells were coated with 10% BSA to prevent sticking. Incubation was carried out in light conditions, approximately 100 µmol/m^2^/s. Wide bore or cut tips were used throughout for handling protoplasts.

### Protoplast Quantification

After the required incubation time (3, 4, 5, 6, and 18 hours for time course experiment, 4-18 hrs for *pDefense-ELuc* reporters) protoplasts were prepared for luminescence measurement. Two-hundred µL of 1x cell lysis buffer (Promega #E3971) was added and samples vortexed and incubated for 15 min. Samples were centrifuged again to collect cell debris and 50 µL of lysate was added to an opaque, flat-bottom, white, solid polystyrene 96 well plate (e.g. Corning® 96 well NBS™ Microplate #CLS3600). Plate was read for total luminescence (Tecan infinite 200Pro, 1000 ms integration 0 ms settle) without addition of substrate, then 50 µL Steady-Glo (Promega #E2520) Luciferin substrate was added, and the plate was read again.

### Protoplast RNAseq sampling

Protoplasts were prepared as described above, using 900 µL protoplast solution (350,000 cells/mL MMG) per replicate, with 4 replicates for each treatment group The *UBI-fLUC* reporter was excluded. Additional replicates of each condition included the reporter to confirm cell-death after 16hrs (Data not shown). At 4 hrs post transfection the protoplasts were pelleted, snap-frozen, and stored at -80 until RNA isolation. Samples from each cultivar were prepared and processed in batches. Qiagen RNeasy Plant Mini Kit was used as directed, (#74904) omitting the first mechanical cell lysis step as freezing and addition of RLT buffer was sufficient to lyse cells. Qiagen RNase-Free DNase Set (#79254) was used as directed. RNA was checked for absence of DNA contamination by PCR and gel electrophoresis and quantitated using Qubit and Nanodrop. Total RNA Illumina sequencing was performed on the NovaSeq platform, in a 2×150bp paired-end configuration, with poly A selection & strand specific library preparation for the 36 samples. The resulting sequence data is deposited in the NCBI Sequence Read Archive (SRA) under BioProject number PRJNA957082.

### RNAseq Analysis

Transcript abundance was quantified with kallisto (Bray *et al*., 2016) using the Ensembl *Triticum aestivum* transcriptome (IWGSC, INSDC Assembly GCA_900519105.1, Jul 2018) plus cDNA sequences for *AvrSr50, Sr50*, *AvrSr35* and *Sr35* as reference. DESeq2 (Love *et al*., 2014) was used to test for differential expression. We defined upregulation at a 4-fold change or greater. GaboSr50 with AvrSr50 was compared against the four control treatment groups; Gabo_AvrSr50, Gabo_EV, GaboSr50_AvrSr35 and GaboSr50_EV. Then, the intersect of the upregulated genes was taken, for a total of 272 genes. For Fielder, Fielder_Sr35_AvrSr35 was compared to the 3 control treatment groups; Fielder_AvrSr35, FielderSr35_AvrSr50, Fielder_Sr35. Again, taking the intersect of the upregulated genes returned 98 upregulated genes. 66 genes were commonly upregulated between both lists (Zenodo Supplemental Data S5). For GO terms analysis, we repeated the comparisons above with lower upregulation of 2-fold change and used the gProfiler webtool and Ensembl *Triticum aestivum* annotation, running the g:GOst program (Raudvere *et al*., 2019). The recommended g:SCS multiple testing correction algorithm was used.

### *pDefense* Promoter Selection

Expression in log fold change and transcripts per million (TPM) was visualized for the 66 commonly upregulated genes, present in both 2-fold upregulated lists used for GO analysis. Thirteen genes were selected based on their overall expression and upregulation in *Avr/R* treatment groups. The entire gene and region were compared in Geneious and 600-800 bp upstream of the 13 genes was selected, including any annotated 5’UTR. These promoter sequences then were synthesized with 5’and 3’ 4bp MoClo promoter overhangs flanked by BsaI recognition sites for Golden Gate cloning as promoter+5’UTR standard parts (Weber *et al*., 2011). Any BpiI and BsaI enzyme recognition sites within the promoter sequence were removed prior to synthesis. Promoters that could not be synthesized were cloned by PCR, using primers (Zenodo Supplemental Data S6) with BpiI recognition sites and 4bp overhang to facilitate assembly into universal level 0 vector *pAGM9121* (Addgene Plasmid #51833). Any BsaI recognition sites were removed as per Grutzner & Marillionnet, (2020).

### *pDefense* Plasmid Constructs

Plasmids were designed in Geneious. The plasmid constructs were designed according to the MoClo syntax, assembled by Golden Gate cloning and verified by whole plasmid sequencing. Supplemental Data (Zenodo Supplemental Data S6) and plasmid maps (Zenodo DOI: 10.5281/zenodo.7844465, link: https://zenodo.org/record/7844465) are available, with all generated plasmids deposited on Addgene.

### *pWDV1* vector

The wheat dwarf virus genomic sequence from the isolate Enkoping1 (GenBank: AJ311031.1) was modified to remove the V1 and V2 genes which encode the viral coat and transfer proteins, respectively. In addition, one and two BsaI type II restriction sites were mutated to enable downstream golden gate cloning steps. All mutations were in the Rep and RepA open reading frames and were silent, with the exception that the mutation generated at amino acid position 263 for RepA results in a Serine to Threonine substitution at amino acid position 235 of the Rep protein which is encoded by an alternately spliced transcript from the same Rep/RepA open reading frame. Two BsaI sites were inserted between the Long and short intergenic regions in place of the removed V1 and V2 genes. The BsaI sites were designed to generate a GGAG overhang downstream of the long intergenic region and a CGCT overhang upstream of the short intergenic region to facilitate level 1 golden gate assemblies as described previously (Engler *et al*., 2014). BsmBI sites were also included at either end of the modified viral genome sequence to facilitate golden gate cloning into the pYTK001 vector (Lee *et al*., 2015) which was chosen for its small size, ease of cloning and to facilitate selection and plasmid replication in bacteria. The modified wheat dwarf virus sequence, called WDV_1, was synthesised as a ‘gene’ fragment by twist bioscience and directly incorporated into the pYTK001 vector by golden gate cloning using BsmBI. We sequence verified plasmids via whole plasmid sequencing. We named the resulting plasmid *pWDV1*. We used *pWDV1* for subsequent cloning of genes by golden gate cloning using the BsaI entry site.

### Ratiometric Assay

All isolation and transfection steps were as above aside from the substitution of the fLUC reporter for the reporter (*UBI-pDefense)* and normalization (*UBI-pRedf*) plasmids. These require the same substrate for activation, but express luciferase with different emission spectra, read by specific green and red filters in the luminescence plate reader.A *UBI-ELuc* construct was also generated to calibrate normalization and deconvolution calculations, necessary to resolve the overlap in emission spectra of the two luciferase reporters (González-Grandío *et al*., 2021). These calibration experiments were performed separately three times and with three replicates per measurement, according to the instructions for Promega Chroma-Glo assay (Chroma-Glo(TM) Luciferase Assay System Technical Manual, TM062). The plate reader was programmed to measure total luciferase, then with green and red filters, with 1000 ms integration time and 0 ms settle time, measured well-wise. Template spreadsheets for performing calculations and a Python script for automating deconvolution and plotting are recorded on protocols.io (https://dx.doi.org/10.17504/protocols.io.q26g7r3zkvwz/v1).

### Cultivar testing

Protoplasts were isolated from nine different wheat cultivars (GaboSr50, Coorong, C6969, Clement, Compair, HeinesPeko, KenyaW1483, AvocetYr8 and Fielder), using the method described above. All were grown in the same conditions as above for 8 days. Where the cultivar carried an *R* gene (GaboSr50, Coorong, C6969) the recognized *Avr* (*AvrSr50*, *AvrSr27* and *AvrSr35* respectively) was transfected alone, with an avr (unrecognized by R protein) as control. In the other six cultivars both the *Avr* and *R* gene were co-transfected. Normalized luminescence was calculated from the ratio of deconvoluted *pDefense-Eluc* to *UBI-pRedf*.

### Multiplex screening

Protoplasts were prepared from cv. GaboSr50 and cv. Gabo wheat leaves and transfected as above. Measurements used the ratiometric reporters and measurements, with *pDefense14-ELuc* as reporter and *UBI-pRedf* for normalisation. The pool of twenty-five *Avr* candidates including *AvrSr50, AvrSr35, AvrSr27, AvrSr22, AvrSr13, AvrStb6, Avr3D1, AvrStb9, AvrPm1a, AvrPm2, AvrPm3d3, AvrPm17*, *AvrSr50-A1, AvrSr50-B6,* and other eleven wheat rust *Avr* candidates used in our studies, including *Pst_104E_10266, Pst_M28_10266, Pst_104E_17331, Pst_M28_17331, Pst_104E_25466, Pst_M28_25466, Pttg_05697, Pttg_27353* and three alternatively spliced version of *Pst_104E_25466.* The pool excluding the *AvrSr50* acted as negative controls, and 1 µg of *pWDV1-AvrSr50* alone acted as a positive control. Data processing and statistics was carried out as above.

## Data availability

All supplemental data is available on Zenodo (DOI: 10.5281/zenodo.7844446., link: https://doi.org/10.5281/zenodo.7844446). All plasmids generated are available from Addgene, with plasmid maps deposited on Zenodo (DOI: 10.5281/zenodo.7844464, link: https://doi.org/10.5281/zenodo.7844464). Code used to perform RNAseq analysis is deposited on Github (https://github.com/ritatam/wheat-protoplast-RNAseq-DESeq2) and all RNAseq data is available on NCBI Sequence Read Archive (SRA) under BioProject number PRJNA957082.

## Acknowledgements

We thank Prof. Robert Park and Prof. Robert McIntosh for providing seeds of cv. C6969, A/Prof. Sambasivam Periyannan for providing seeds of cv. Fielder, cv. Gabo, cv. GaboSr50, and cv. Coorong, A/Prof. Peng Zhang for providing seeds of Federation*4/Ulka, Dr. Michael Ayliffe and Dr. Peter Dodds for providing *pUBI-AvrSr35*, *pUBI-Sr35*, *pUBI-AvrSr50*, and *pUBI-Sr50*. We are grateful to Dr. Cecile Lorrain for useful suggestions on AvrStb6/Stb6 interaction. We thank Jasper Lee for plant maintenance. B.S. acknowledges the financial support by the Australian Research Council via a Future Fellowship (FT180100024). This work was supported by computational resources provided by the Australian Government through the National Computational Infrastructure (NCI) under the ANU Merit Allocation Scheme.

## Author Contributions

S. W., B. D., J. P. R., and B. S. conceived the project. S. W., B. D., R. T., L. M., N. H., S. S. K., E. C., and J. G. performed the experiments, S. W., B. D., and B. S. wrote the manuscript. R. T., E. C., and J. P. R. provided extensive feedback on the manuscript. B. S. secured funding. All authors read and approved the manuscript.

## Supplemental Material Legends

**Supplemental Figure 1: Diagram of plasmids used in protoplast assay. (A)** *pDefense14* consists of D14 promoter that drives expression of the green-shifted E-Luc cassette. *pDs 1-7, 10, 11, 12, 15 and 16* were constructed with the same modular plasmid components, replacing the relevant defence-induced promoter **(B)** UBI-*pRedf*, used for normalization, consists of a red-shifted coding sequence driven by *UBI* promoter. **(C)** Example *Avr* candidate plasmid, with *pWDV1* self-replicating plasmid backbone.

**Supplemental Figure 2: Preliminary successful screening of *pDefense2, -14* and *-15* defense reporters.** Normalized luminescence for *pDefense14,* (panel **A**)*, pDefense 15* (panel **B**) and *pDefense2* (panel **C**), in three wheat cultivars. Left to right, cv. C6969 which carries *Sr35*, cv. GaboSr50 which carries *Sr50* and cv. Coorong, which carries *Sr27*. *Avr+R* treatments were transfected with plasmid which expressed a recognized *Avr, avr+R* treatments expressed a non-recognized *avr* and *R* alone expressed an empty vector. Reporters were normalized against *UBI-pRedf* for each replicate. The three reporters produced an increase in normalized luminescence in treatments with Avr/R interaction, across the three *Avr/R* pairs tested.

**Supplemental Figure 3: *pDefence* reporters confirm *AvrSr50* variants evade recognition by a single amino acid substitution.** *AvrSr50* variants; avrsr50-A1 (Q121K), avrsr50-B6 (avrsr50QCMJC) and AvrSr50-C (AvrSr50RKQQC), from Ortiz et al., (2022) were synthesized and cloned into *pWDV1* vectors. avrsr50-A1 had a single amino acid substitution (Q121K) that prevented recognition by Sr50, similar to avrsr50-B6, while AvrSr50-C had six amino acid changes but was still recognized by Sr50. One-way analysis of variance (ANOVA) and post hoc Tukey’s HSD test was used to assess differences among means of normalized and red-filtered luminescence measurements for each treatment. Treatments marked with common letters were not significantly different (P >0.05).

**Supplemental Figure 4: The *Stb6* gene amplification profile in ten different wheat cultivars.** The *Stb6* (exp: 782 bp), and wheat Polymerase A1 gene (*TaPolA1*; exp: 500 bp) partial amplifications were performed. PCR products of 3 µL were loaded on 1% agarose gel. M: 1 kb Plus DNA ladder (NEB #N3200).

**Supplemental Figure 5: *pDefense* reporter can detect lower purity *Avr* plasmid.** In **(A)** both maxi prepped (left) and mini prepped (right) *AvrSr35* plasmid was co-transfected with *R* and *avr* plasmids, in protoplasts from cv. Fielder. **(B)** shows corresponding non-normalized red-filtered luminescence. One-way analysis of variance (ANOVA) and post hoc Tukey’s HSD test was used to assess differences among means of normalized and red-filtered luminescence measurements for each treatment. Treatments marked with common letters were not significantly different (P >0.05).

**Supplemental Material S1:** Luminescence data for Figure 1, protoplast time course experiment.

**Supplemental Material S2:** List of samples and treatment groups for RNAseq experiment.

**Supplemental Material S3:** Lists of Gabo/GaboSr50, Fielder and all cultivar genes upregulated by 2-fold.

**Supplemental Material S4:** GO terms analysis Query tables and g:profiler results tables

**Supplemental Material S5:** Lists of Gabo/GaboSr50, Fielder and all cultivar genes upregulated 4-fold.

**Supplemental Material S6:** List of primers used.

**Supplemental Material S7:** List of plasmids used.

**Supplemental Material S8:** Luminescence data used for generating Figures 3-7.

**Supplemental Material S9:** Statistical output for Figures 3-7.

**Supplemental Material S10:** Data and statistical output for Supplemental Figures 2, 3 and 4.

