## Supplementary figures and images for "Multiplexed effector screening for recognition by endogenous resistance genes using positive defense reporters in wheat protoplasts"

### Supplemental Figures

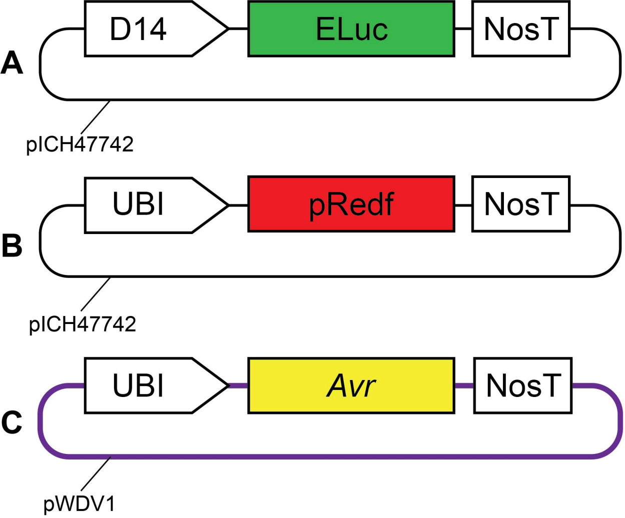

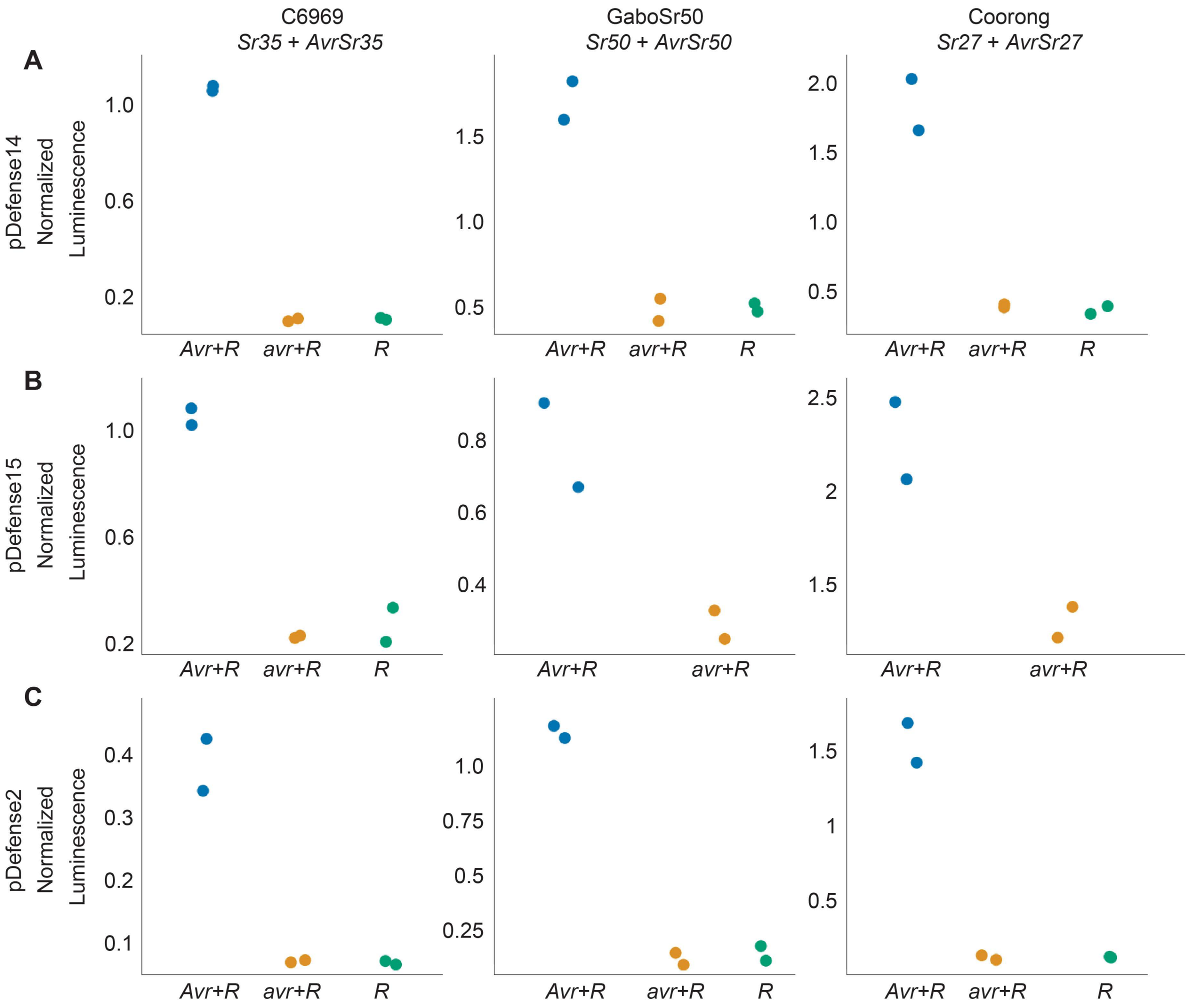

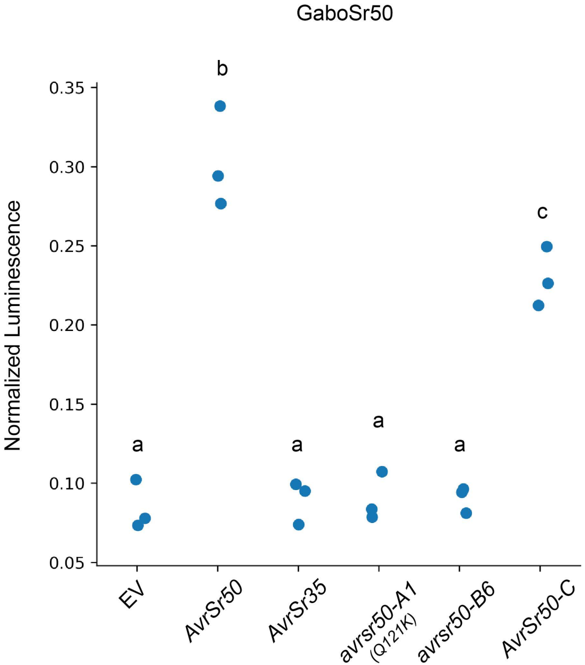

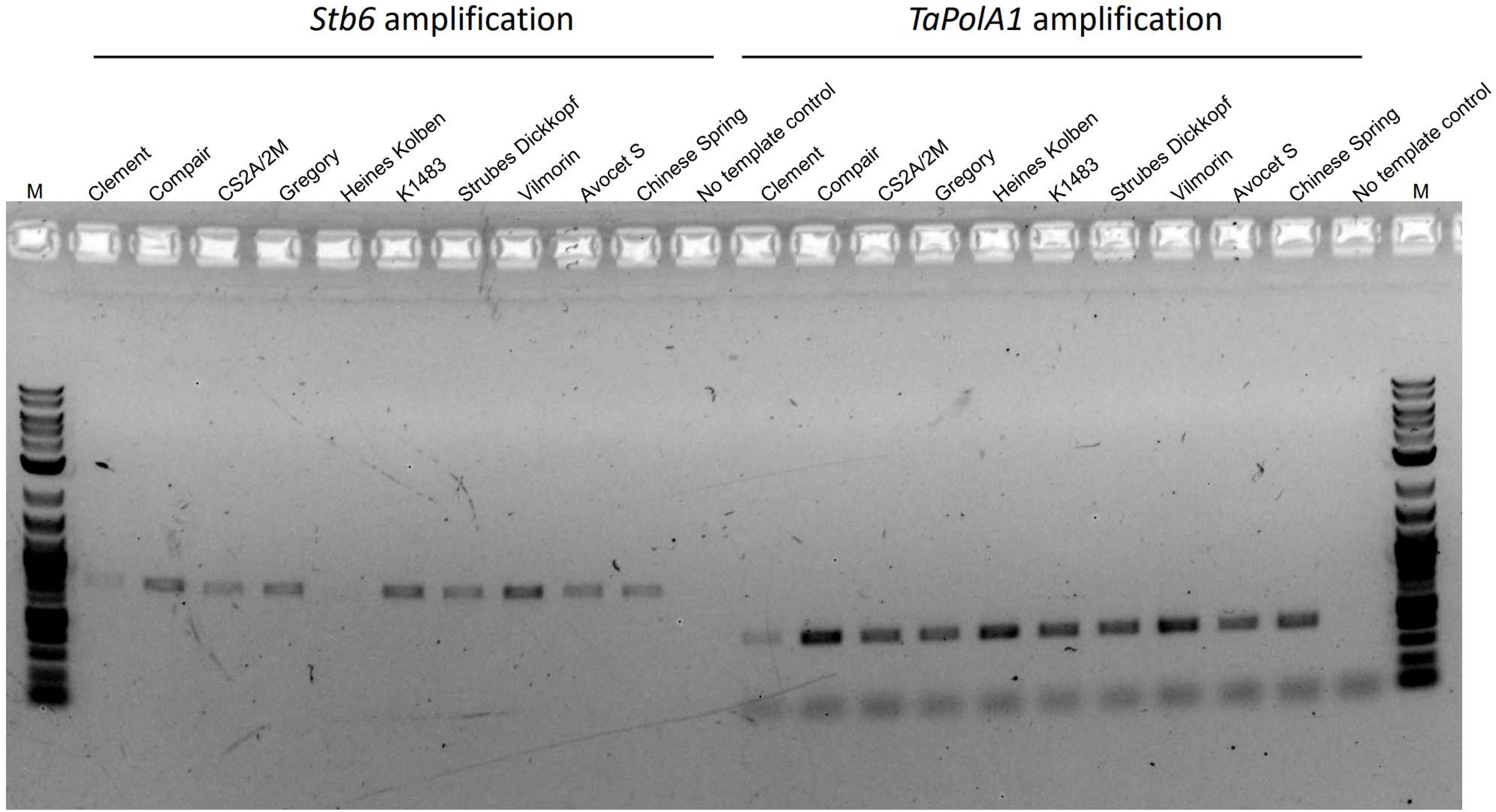

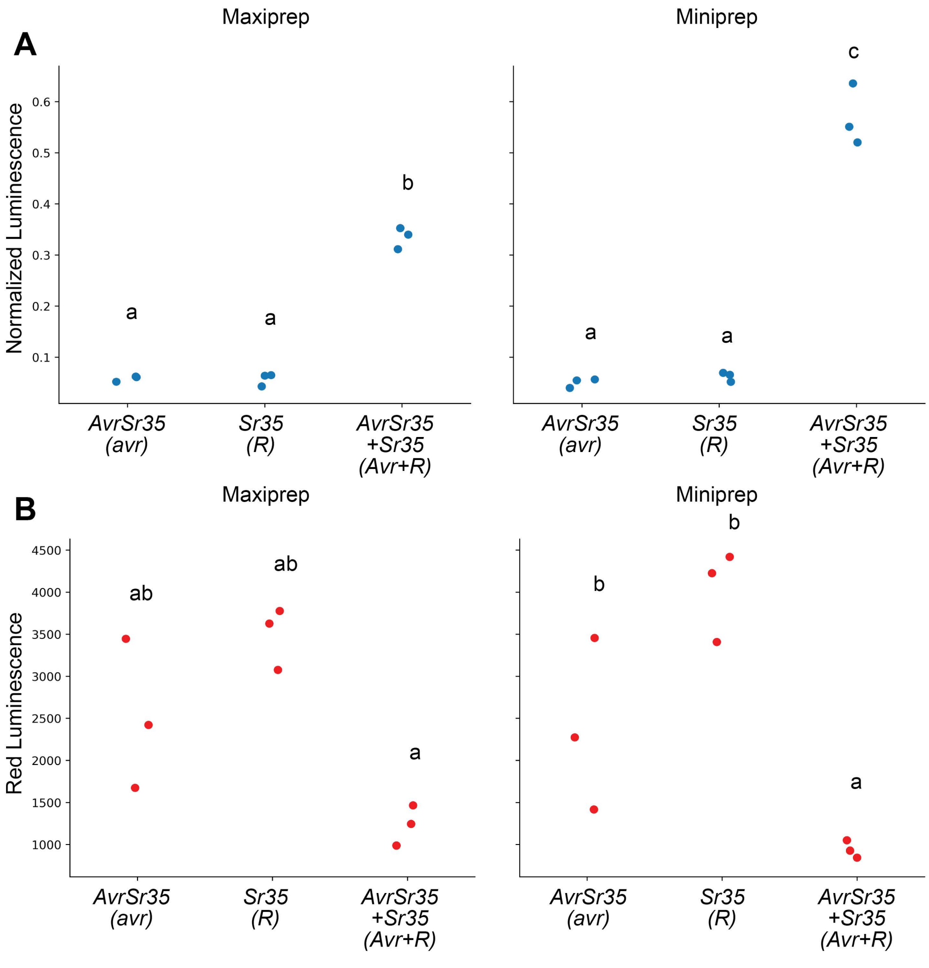
